# Study buddy learning is associated with academic success in undergraduate science courses

**DOI:** 10.1101/2025.05.20.655056

**Authors:** Achol Jones, Gabrielle Reznik, Al Rohet Hossain, Veronica Dudarev, Bowen Hui, Patrice Belleville, Warren Williams, Eden Fussner-Dupas, James Enns

## Abstract

This study reports on the outcomes of a formal peer-learning program for undergraduate science students. We implemented a cohort-based study buddy program in which high-scoring students (mentors) and lower-scoring peers (protégés) were invited to form study groups based on their first assessment scores in a given course. We then compared the performance of participants versus non-participants on subsequent assessments, while examining separately the effects for mentors and protégés. Results showed similar participation benefits for students in both mentor and protégé roles. These findings emphasize the value of reducing barriers to collaborative learning in higher education and highlight the study buddy program as a cost-effective and scalable approach to enhance students’ educational experiences.

## Introduction

Learning-by-teaching is known to boost student learning, their retention of material, and socio-emotional engagement with their peers [1–10]. In higher education, instructors have first-hand experience of how this approach can improve comprehension of course material. When an educator breaks down a concept to make it easier for students to grasp, both the student and the instructor learn the material more thoroughly, even though the instructor is already an expert in the field. Here we report on our implementation of a cohort-based peer-learning program targeted broadly for undergraduate university science courses. In our program, volunteering students in the same course were grouped into small teams of two to four, based on their scores on a first assessment in the class. Students who scored above the mean on the first assessment (hereafter referred to as *mentors*) were paired randomly with students who scored below the mean (*protégés*). Pairing students in this deliberate way allowed us not only to compare the scores of participants versus non-participants on subsequent assessments, but also to examine the benefits separately for mentors and protégés.

### Collaborative Cognition in the Lab

Cognitive teamwork has previously been studied extensively in laboratory settings. A meta-analysis of 26 studies comparing group behavior in a wide range of cognitive tasks (i.e., recall of pictures and words, visual search for targets, problem solving, model building, reasoning) pointed to three critical variables for predicting better performance [11]. First, groups composed of friends outperformed those composed of acquaintances or strangers across all task types. Second, the benefit of friendship was greater as tasks became more complex, meaning there were a larger number of task elements and outputs to coordinate [see also 12]. Third, the benefits of friendship increased along with group size.

Laboratory studies have also pointed to importance of measuring group performance in such a way that it distinguishes between mere statistical facilitation and social interaction. To appreciate this distinction, consider the likelihood that two tossed coins will each land ‘heads’ under two conditions: (1) the coins are tossed in separate rooms, (2) the coins are tossed in the same room while attached with sticky tape. In one study [13], pairs of participants performed a difficult speeded visual search task under these analogous conditions. The main finding was that the size of the social interaction effect (i.e., beyond that of statistical facilitation) was correlated with the strength of the pre-existing friendship present in the teams. In a follow-up study [14], the same authors demonstrated this friendship advantage in task performance, in addition to verbal communication, was reliant on body language cues. The ‘sticky tape’ connecting the teams’ individual performances appeared to be related to the fluency of their social exchanges.

In summary, laboratory studies have pointed to the importance of (1) friendship in creating effective teamwork, (2) comparing team performance against a standard that controls for the mere statistical facilitation that occurs when the performance of two or more persons is compared to a single person, and (3) that the benefits of friendship in teamwork likely includes mechanisms ranging from increased motivation, to more fluent communication, and to communication that is not only verbal, but includes visible body language. But what these studies do not address is the impact that collaborative cognition may have on learning in a non-controlled setting, such as performance in an undergraduate science course.

### Collaborative Cognition in the Wild

In contrast to the many studies of collaborative cognition in the lab, there have been fewer studies of teamwork in real world situations. It is important to bear in mind that these real-world studies necessarily lack many of the controls one expects to see in a laboratory study. Yet, this limitation in being able to make strong causal statements at the conclusion of a study can be offset by the benefits that come from studying human behaviour ‘in the wild.’ After all, what benefits do strong causal statements provide, without some understanding of whether the same phenomena occur in the natural world and by documenting the conditions under which they occur?

Previous studies within educational contexts have shown that students learn best when they split their time between the roles of learner and teacher, following the 50/50 Feynman principle [15,16]. Other studies have shown that students make a greater effort to learn with the intention to teach a teaching agent, than they do for themselves—a demonstration of the phenomenon called the protégé effect [17]. A few institutions have implemented cohort-based peer-learning, though these are found mostly in professional degree programs with standard schedules [9, 10]. Many colleges and universities support formal mentorship programs, where senior and junior scholars are paired in peer-tutoring setups, to offer extra support to students [1–4]. Yet, many students in higher education continue to work alone, often relying on solo efforts, possibly because they perceive a zero-sum competition for class rankings. This isolation was made worse by the physical distancing measures required due to the COVID-19 pandemic [18,19].

We launched our study buddy program in undergraduate science courses during the COVID-19 pandemic to help reduce barriers to forming study partnerships. We hoped that pairing students in the same class would drive them to engage more deeply with the material, regardless of their initial comfort with the subject matter. We hoped that these pairings would lead in many cases to the kinds of friendship benefits documented in laboratory studies. We also believed that learning-by-teaching was an underused, cost-effective way in which to catalyze both students’ social networks and their mastery of course material. Finally, we designed our study so that we could directly compare the performance of students in teams with students of equivalent potential, based on an initial assessment in each course, that were not participating in teams. As our results show, encouraging study buddy teams proved to be a powerful way to benefit both struggling students and top achievers, helping them all get more out of their studies.

## Method

Participants include 1,353 students at the University of British Columbia (UBC), a government-funded research-intensive Canadian university: 484 students over four terms in a second-year Neuroscience Statistics course (2021-2024), 620 students over two terms in a second-year Biochemistry course (2023-2024) and 249 students in one term of a third-year Biochemistry course (2024). Because some students did not complete all the assessments the final sample size was 1,314 students. After students had received feedback on their performance in the first exam-based assessment of the term, typically about one month into the course, they were invited to participate in the study buddy program by a member of the research team.

The initial version of this program began informally in a single course, the Neuroscience Statistics course in 2021 and 2022. The instructor informed students after releasing grades on the first assessment in the course that occurred during the fourth week of classes (February 4, 2021). Students were told verbally that, to encourage groups studying, they could volunteer in an informal study buddy program, by sending an email to the instructor, who would match students in pairs, on a first-come-first served basis, with one member having scored below the average on the first course assessment and one member having scored above average. These volunteer emails were returned with a brief welcome message, along with the email of the matched partner. Students were informed in class that any analyses of their subsequent performances in the course would be done on an aggregate basis, without any reference to personal identifying information other than their participation as study buddy or solo studying students. These data were accessed for evaluation purposes of the program following the final exam on April 27, 2021. Personal identifying information was replaced with random numerical codes in these files. Article 2.5 of TCPS2, the Canadian policy framework governing research ethics, indicates that quality assurance activities (e.g., testing the effectiveness of a pedagogical technique) do not require institutional research ethical review.

These procedures were formalized in June 2023, when the study team applied to the UBC Board of Research Ethics to conduct study buddy programs in several Biochemistry courses, in addition to the Neuroscience Statistics course. Approval of all procedures was granted on June 19, 2023 and then renewed again on May 23, 2024. In the initial recruitment survey, September 15, 2023, students were asked for basic contact information, meeting preference (online, weekdays, evenings & weekends), and preferred role (mentor, protégé, or either). Note that these additional parameters were used only for high-level guidance in matching categorization. Beyond this initial categorization, students were matched randomly, pairing students with scores above and below the mean. Once matches were complete, students were contacted by email stating that they are participating in the program and that it was their responsibility to get in contact with their buddies (roles were not assigned to individuals in this communication). Informed written consent was given by all participating students prior to participating in the survey, both those who volunteered to participate in the study buddy program as well as those students who did not participate in formal study groups, but consented to have their grades used as comparison data in the analyses of the program’s success. Data collection ended for the data reported in this study on April 30, 2024, and some team members (those teaching each class) had access to the identifying information prior to it being anonymized for analysis by other team members. Additional method details are given in the Supporting Information, including the number of participating students in each course and descriptive statistics of the assessments in each course (Table S1).

Students participated in the program at an average rate of 30%, with a range from 18% to 50% across the seven sections of three courses. Without any further intervention, we then monitored the academic progress of the students in the study buddy teams over subsequent assessments in the course, comparing them to non-participating students within the same course. We will refer to the latter as solo students, while acknowledging the possibility that some students studied with friend(s) without participating in the study buddy program.

## Results

The data and code used in the analyses reported in this study are available at https://osf.io/85zhf/?view_only=34ece155ffb34729888461d28647e926. Note that all identifying features of students in these courses have been removed from the data set. Here we report the aggregated highlights of our findings; a more comprehensive report is given in the Supporting Information. We began by asking whether students who participated in the program, differed in their first assessment scores from non-participating students, by comparing their respective distributions of first assessment scores (Figure S1). Both distributions had almost identical means and standard deviations. We interpret this to imply that participating students were not differentially motivated in their academic pursuits, at least by virtue of their scores on the first assessment.

Assessment grades in each course were converted to z-scores to standardize the means and variance across courses. For some analyses, change scores were also computed for each participant, consisting of the difference in z-scores between the first and last assessments in a course. To express costs and benefits of the changes in practical terms, z-scores and change scores were then converted to grade percentages by multiplying them by the standard deviations for each assessment. (Figures S2 and S3). These data were analyzed with analysis of variance (ANOVA) using R [20].

The aggregated mean-centered results, across all courses in the dataset, are shown in Figure 1A. Students participating in the program gained an average benefit of more than 2.0%, when compared to non-participants, whose grades declined slightly on average from the first to last assessment. Team scores were on average significantly higher than solo scores, F(1,1258)=4.29, p=0.039, and the interaction of role x assessment was statistically significant, F(1,1258)=13.69, p<0.001. We observe the same trends in study buddy benefit when all assessments in each of the participating courses are considered (Figure S2).

**Figure 1.**
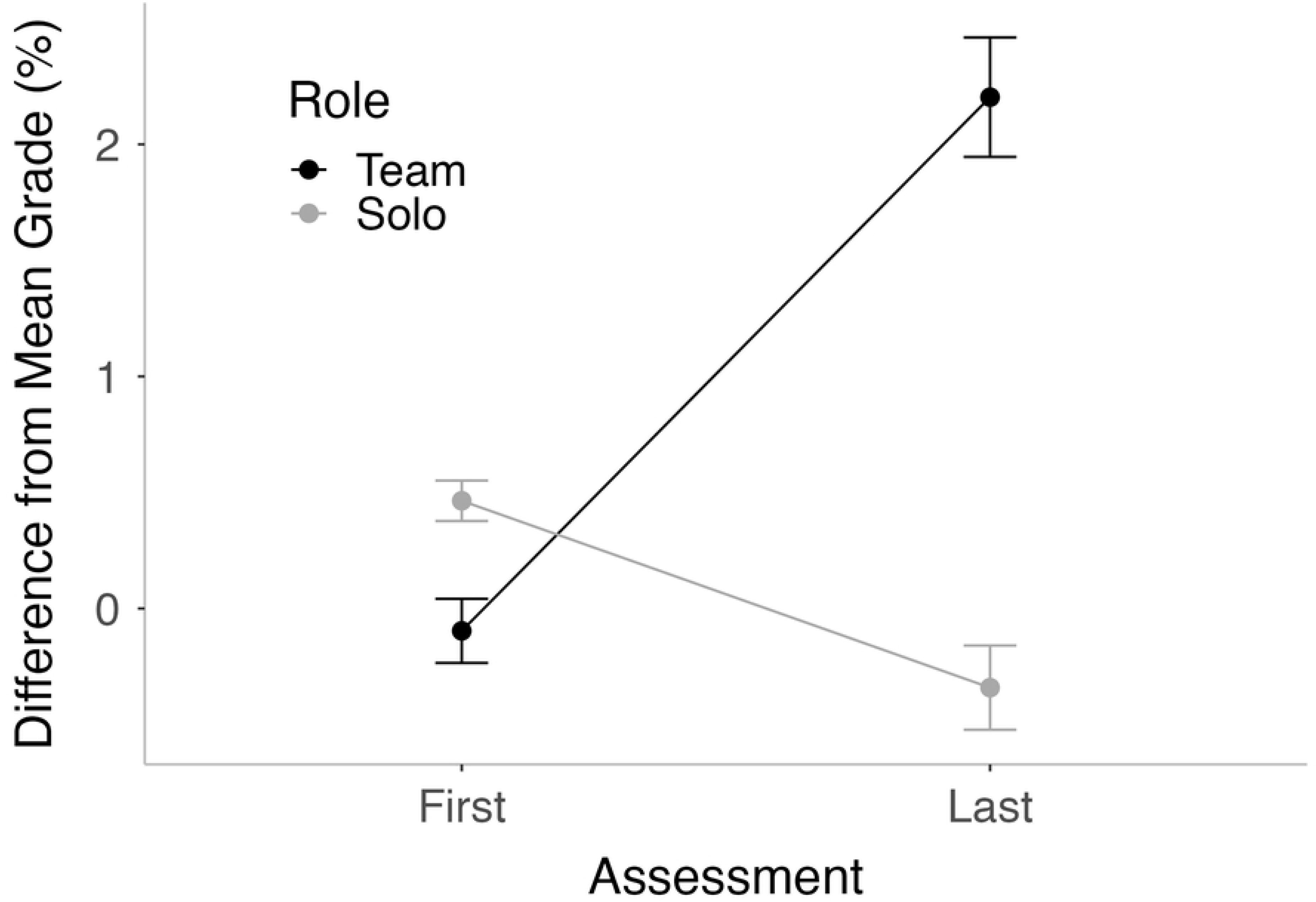
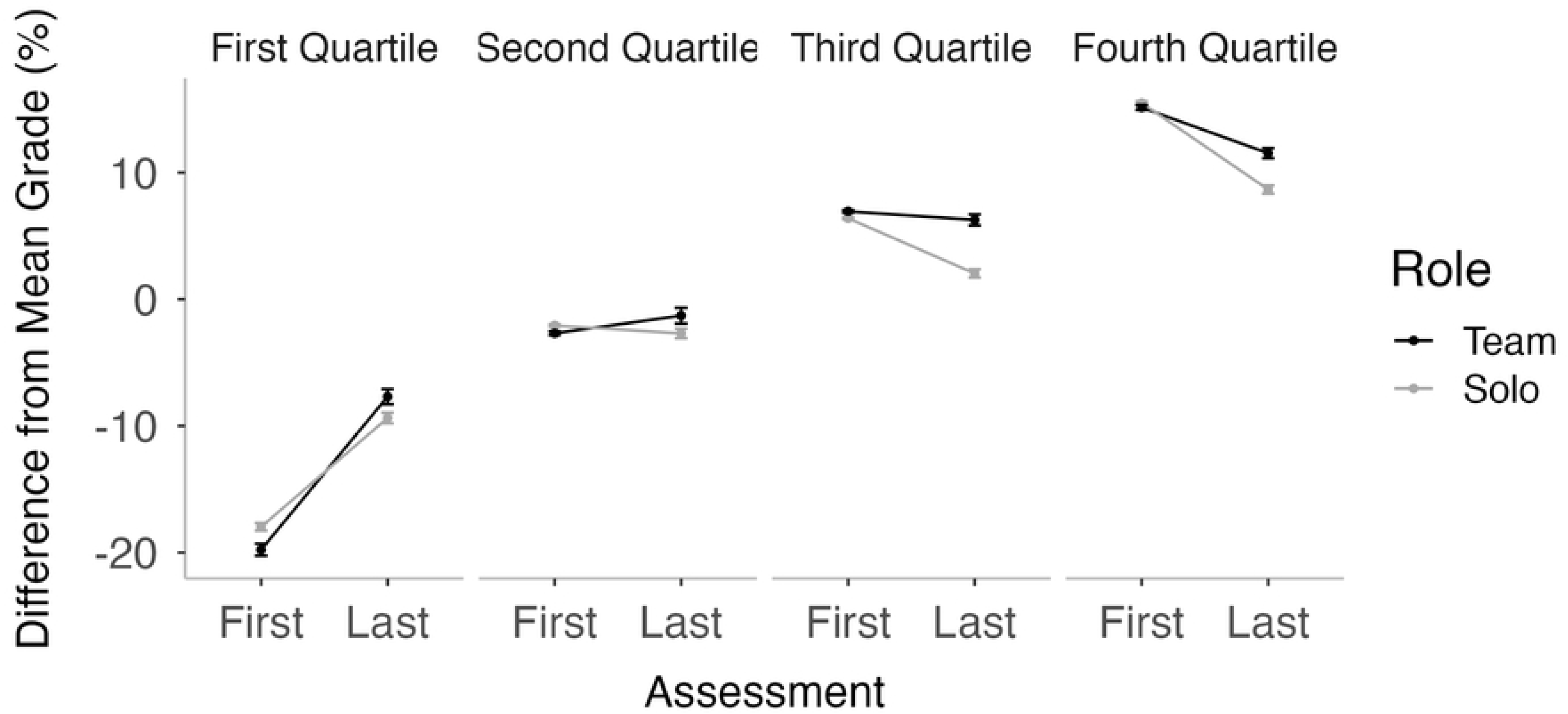
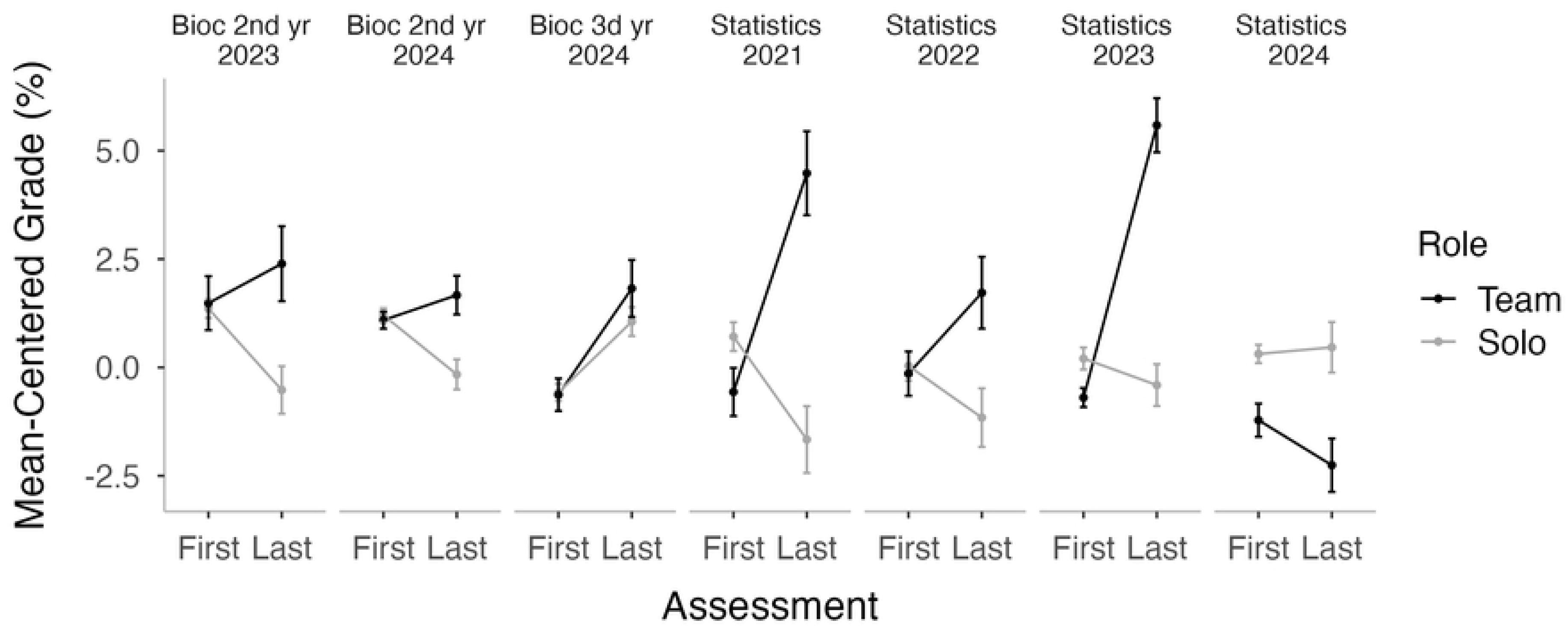
**(A)** Mean-centered scores on the first and last assessments in a course, aggregated results across all courses in the dataset. (B) The same data shown separately for each quartile of scores on the first assessment. (C) The same data shown separately for each course in the dataset. Errors bars are +/-1 standard error.

We then examined these data in greater detail by dividing student’s scores on the first assessment into quartiles (Figure 1B). We refer to students in the lower two quartiles as ‘protégés’ and those in the upper two quartiles as ‘mentors,’ to reflect their potential roles in a study buddy program. The analyses showed that the benefits of team studying were similar for mentors versus proteges, as indicated by the non-significant interaction of assessment x role x quartile, F(1,1258)=0.20, p=0.897. This finding indicates that solo students with mentor potential (those who scored above the mean on the first assessment) had lower scores on average than team students, and that solo students with protégé potential (those who scored below the mean on the first assessment) also had lower scores on average than team students. These results likely underestimate the benefits of participating in the program, as some solo students may have not worked solely on their own and some team students may have not engaged with their partners.

We also analyzed these data to look at the results for each course separately (Figure 1C). There were no measurable differences between the outcomes in these courses when considered in the context of the overall analysis: the interaction of assessment x role x course was not significant at the .05 level, F(1,1258)=1.85, p=0.086. However, when we examined the change from first to last score in each course separately, four of the courses showed significant differences between team and solo students, with an average difference of more than 3.0%. The three courses not showing significant differences had an average difference of less than 1.0%. We note that a team benefit of 3% amounts in many cases to a full grade letter change in the grading scheme for students at this institution. We also note that these are likely conservative estimates of the effects, considering that we do not know how much time students actually spent together, how their time was spent, nor their level of commitment to the program, or if non-enrolled participants were also studying in small groups with other classmates. Despite these obvious limitations, these data demonstrate the efficacy of the program on a population level for protégés and mentors alike. Parallel analyses, examining all the assessments in each course, are given in Supporting Information (Figure S4 and Table 2).

In a secondary analysis — involving only a subset of the data — we tested for possible influences in the average change scores related to team heterogeneity in first assessment scores. There is a mixture of existing evidence concerning what makes an optimal team in a peer-learning context to maximize student engagement and success. Some studies report that students in study groups with a heterogeneous composition perform better and rate their experiences as more satisfying [21]. Other educators advocate for matching students with other learners who have achieved similar academic success, under the rationale that this will encourage more fluid and equitable sharing of information within the team. Our program design of randomly linking mentors and protégés allowed us to look for empirical evidence for this issue in the data from 2023-2024, where we kept detailed records of team composition for the first time. We began by matching each of the 310 team students in this data set with a random solo student receiving the same score on the first assessment in the same course, to provide a matched baseline against which team benefits could be compared. This gave us a total data set of 620 students, with half of them studying in teams and the others solo. In both groups, students with assessment scores above the mean were labeled ‘mentors’ and those with scores below the mean ‘protégés’. We coded for team heterogeneity by using the standard deviation of first assessment scores in each team, and then binning these scores into four quartiles.

The average change scores from the first to last assessment, as a function of team heterogeneity is shown in Figure 2, separately for students scoring above versus below the mean on the first assessment, and separately for participating (team) vs non-participating (solo) students. Consistent with our aggregated data, we observe that team students had significantly larger percentage change scores overall than solo students, F(1,301)=24.62, p < .001. But beyond the overall team benefit seen in the main effect, there was a different pattern for students scoring above versus below the mean on the first assessment: the interaction of team heterogeneity x first assessment score was significant, F(3,301)=7.84, p < .001. When we focused on the students with scores above the mean (left panel in Figure 2), other than the significant benefit of team participation overall, F(1,177)=6.15, p < .014, there was no interaction of team heterogeneity x student role, F(3,177)=0.36, p=.782, reflecting that the benefit of team participation was similar at all levels of team heterogeneity. This means that mentors received a similar boost from learning-through-teaching, regardless of whether they were assisting students with lower or more similar grades to their own. A similar pattern emerged for students scoring below the mean on the first assessment (right panel in Figure 2).

**Figure 2.**
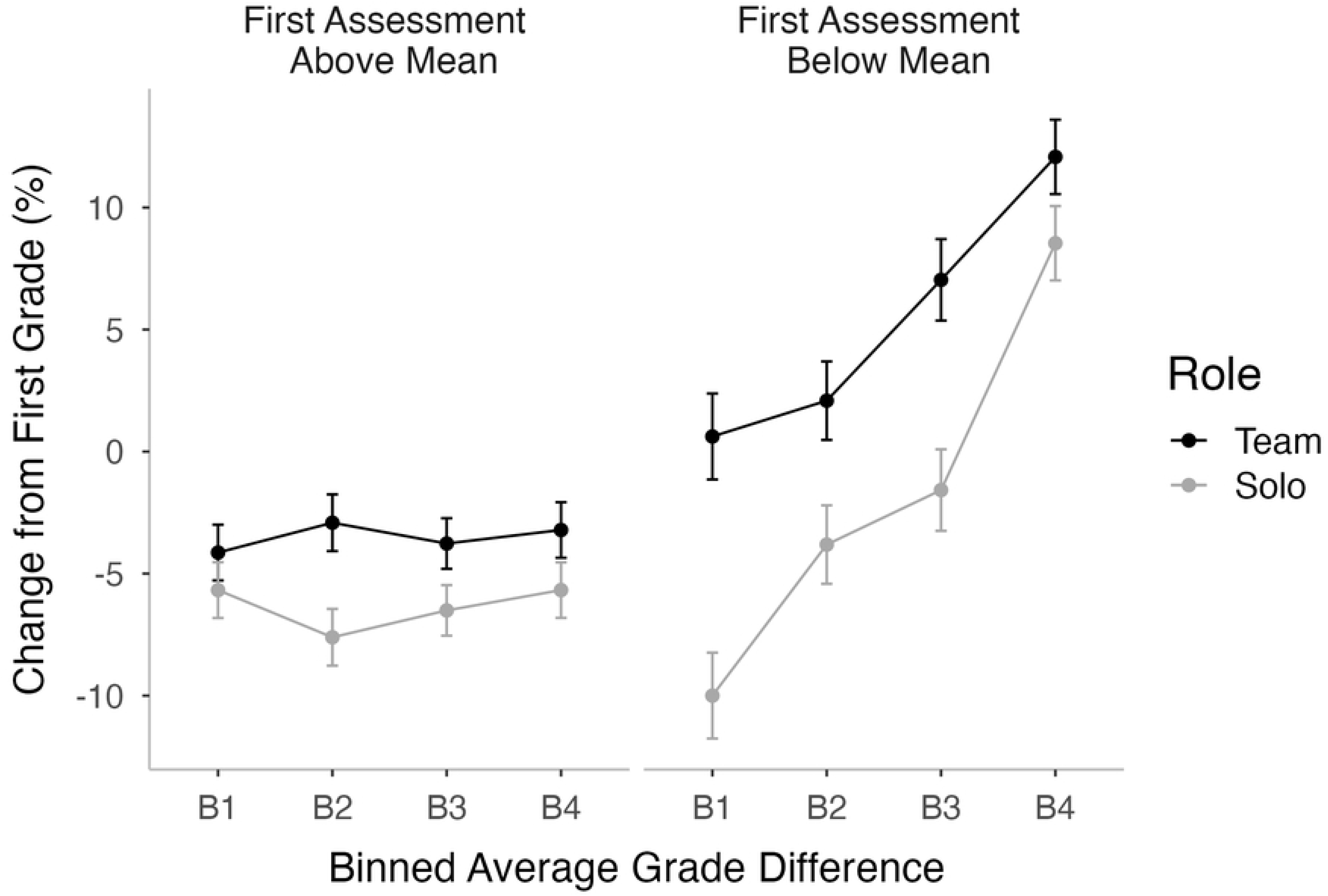
Mean percentage change in grade following the first assessment, as a function of average grade differences among team members, student role (team, solo), and grade on the first assessment (above mean, below mean). Errors bars are +/-1 standard error.

For these students, change scores generally rose as team heterogeneity increased, F(3,124)=10.04, p < .001, reflecting the fact that students with low scores on the first assessment tended to do better overall on subsequent assessments, regardless of their participation. However, the benefits of team participation, F(1,124)=16.64, p < .001, showed no interaction with team heterogeneity, F(3,124)=0.76, p=.516. As with mentors, the benefit of team participation for proteges was evident at all levels of team heterogeneity. Parallel analyses, examining all the assessments in each course, are given in the Supporting Information (Figure S5).

## Conclusion

This study provides empirical support for the claim that undergraduate science students benefit academically when they are encouraged to work together in a formalized study buddy program. Although the size of the benefits varied from course to course, and from student to student, the most robust finding we report is that the benefits at a population level were similar in size for protégés and mentors alike. Note that the finding of similar benefits for these two student roles need not imply that the mechanisms are the same. It is possible that the benefits for students initially scoring below the mean on a first assessment may derive benefit from being tutored by a peer, whereas the benefits to mentors may derive in part from the learning-by-teaching effect [1–10]. Moreover, students in both roles may be benefiting from the implicit nudges of the program to distribute their learning over time rather than cramming before assessments [22]. Future research targeted more directly at these factors will be needed to assess their relative contributions.

In the laboratory studies of collaborative learning, clear links between friendship, and modes of communication enhance a team’s ability for problem solving. Are we observing the same phenomena in this in-the-wild study looking at student behavior as reflected in academic outcomes in science courses? Teams clearly are performing better then students who elected to study alone. We are also observing the enhanced achievement —beyond the sum of the parts — which previous laboratory studies have reported [13,14]. It is notable that we observed a benefit for both the mentors and the protégé, which mirrors the success of teams who had established friendships prior to the laboratory task studies. In future studies we will explore what, if any, impact the study buddy program had on fostering friendships. Is friendship or relationship building a cornerstone of learning, and of mastery, in higher education classrooms? And as an extension, what impact does the size of the study group size and how learners are engaged in study buddy partnerships impact outcomes?

A central pillar of higher education is providing enriching opportunities for undergraduate scholars to learn and engage with course materials, so they can retain and apply their discipline-focused understanding in their future careers. There is now a noticeable shift in appreciation of the overall approach to teaching in higher education, where classrooms are being transformed into learning ecosystems with teachers and learners laterally distributing knowledge and contributing to the culture of learning. Yet, barriers remain in actively building these frameworks where students can study together and support each other’s learning. We offer this report in the hope that the initial success of our study buddy program will inspire instructors and curriculum developers to continue removing the barriers to collaborative learning. We also wish to emphasize that our implementation of instructor-facilitated student study partnerships is a cost-effective and scalable means of enhancing the quality of learning in higher education classrooms, requiring minimal additional intervention from instructors and teaching assistants.

## Supporting Information

**Table S1**. **Average grades in course assessments.** The number of students participating by course, year, study role (team, solo), and the average assessment grades (standard deviations in parentheses).

**Figure S1.**
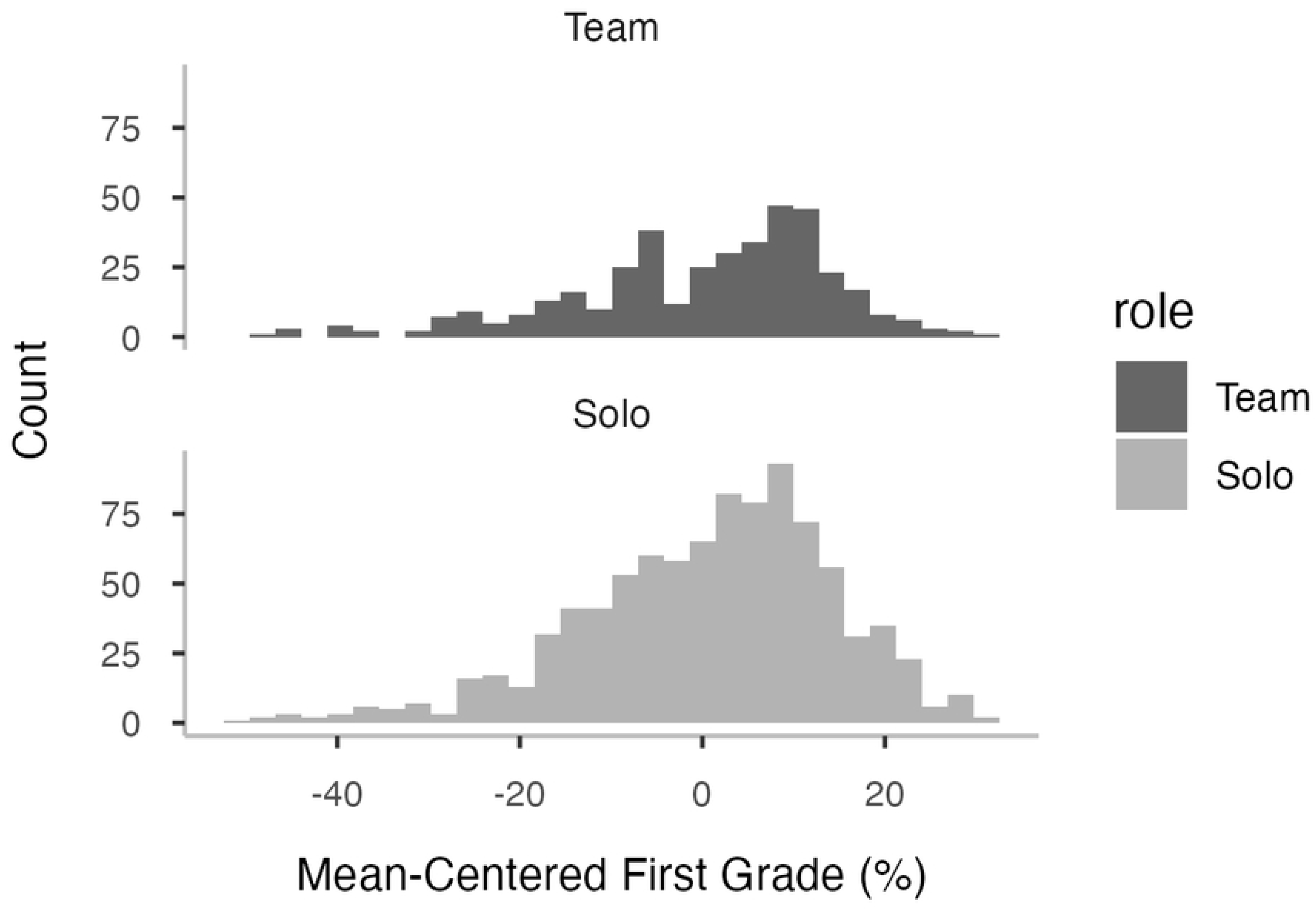
A comparison of first assessment scores for participating and non-participating students. The distribution of scores for the first assessment in each course, separately for 397 students participating as study buddy members (team) and the 917 students studying alone (solo).

**Figure S2.**
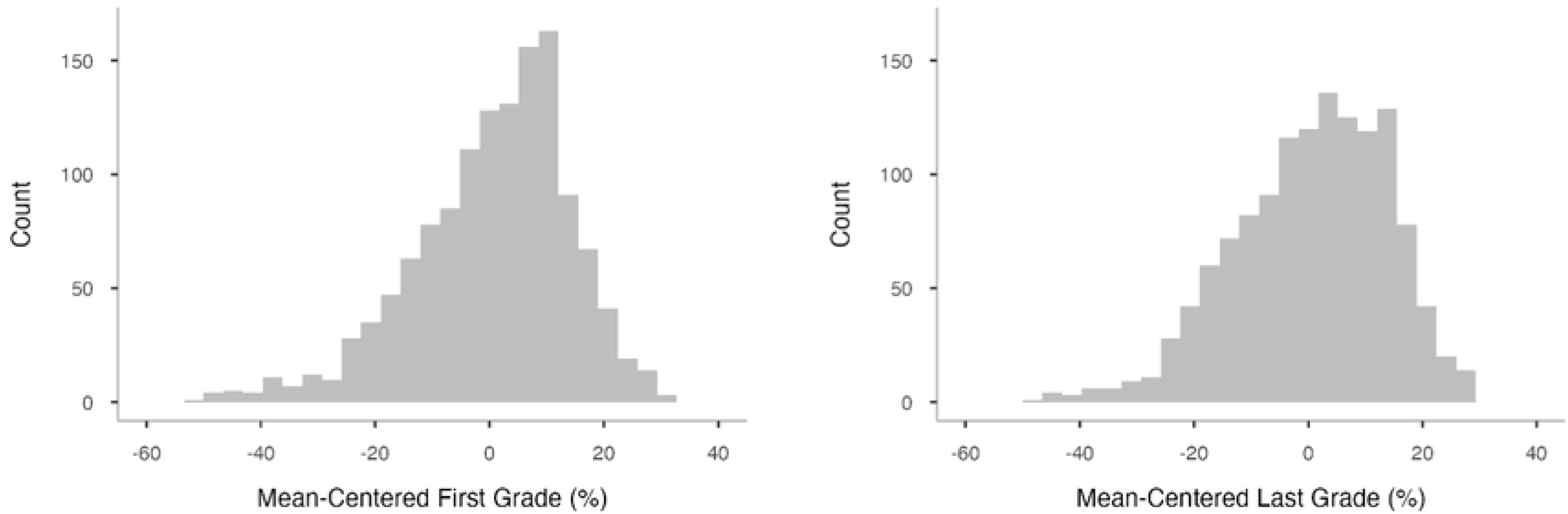
Grade Distributions in the Final Exam Approach. The combined distribution of grades, expressed as differences in percentage scores from the mean, for first and last assessments across all courses.

**Figure S3.**
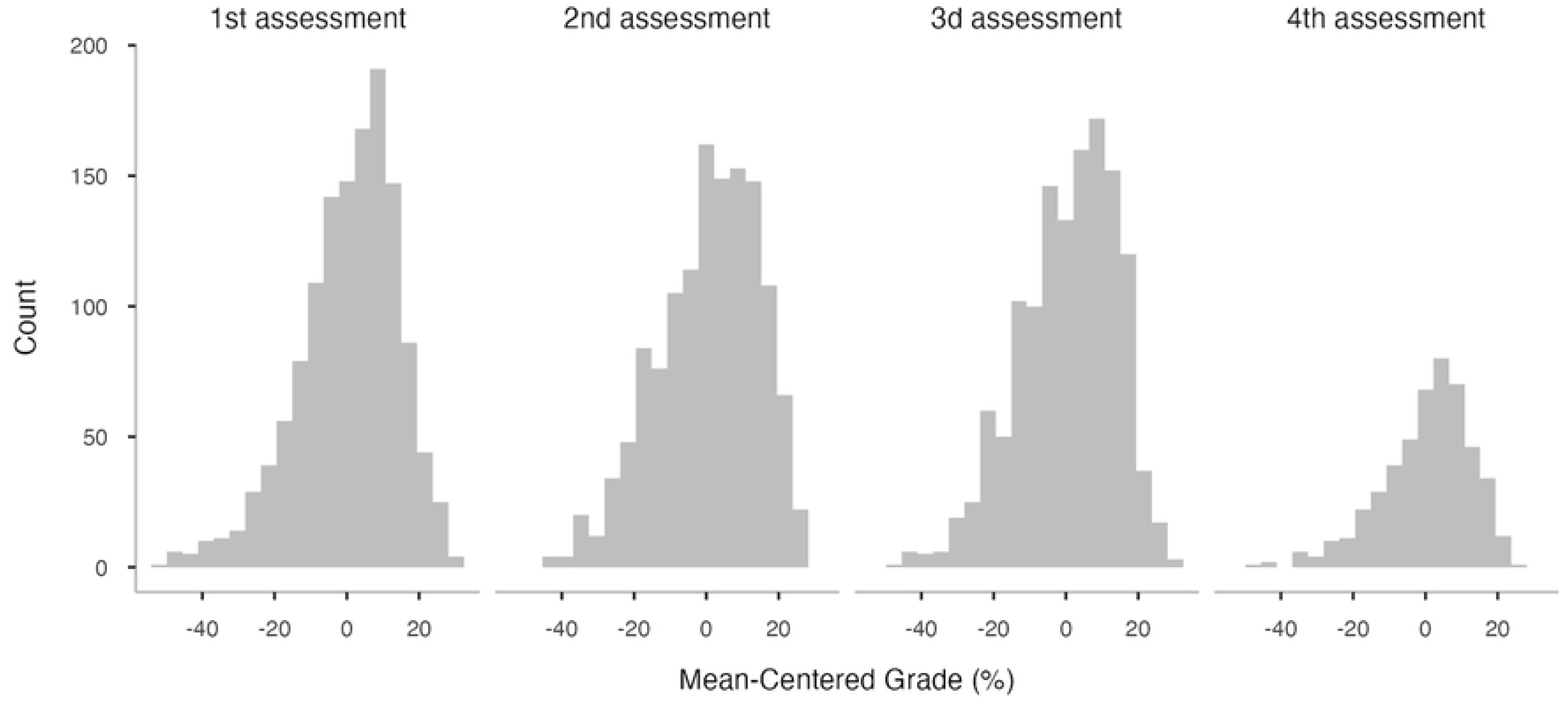
Grade Distributions in the Total Change Approach. The combined distribution of grades, expressed as differences in percentage scores from the mean, for all assessments in each course.

**Figure S4.**
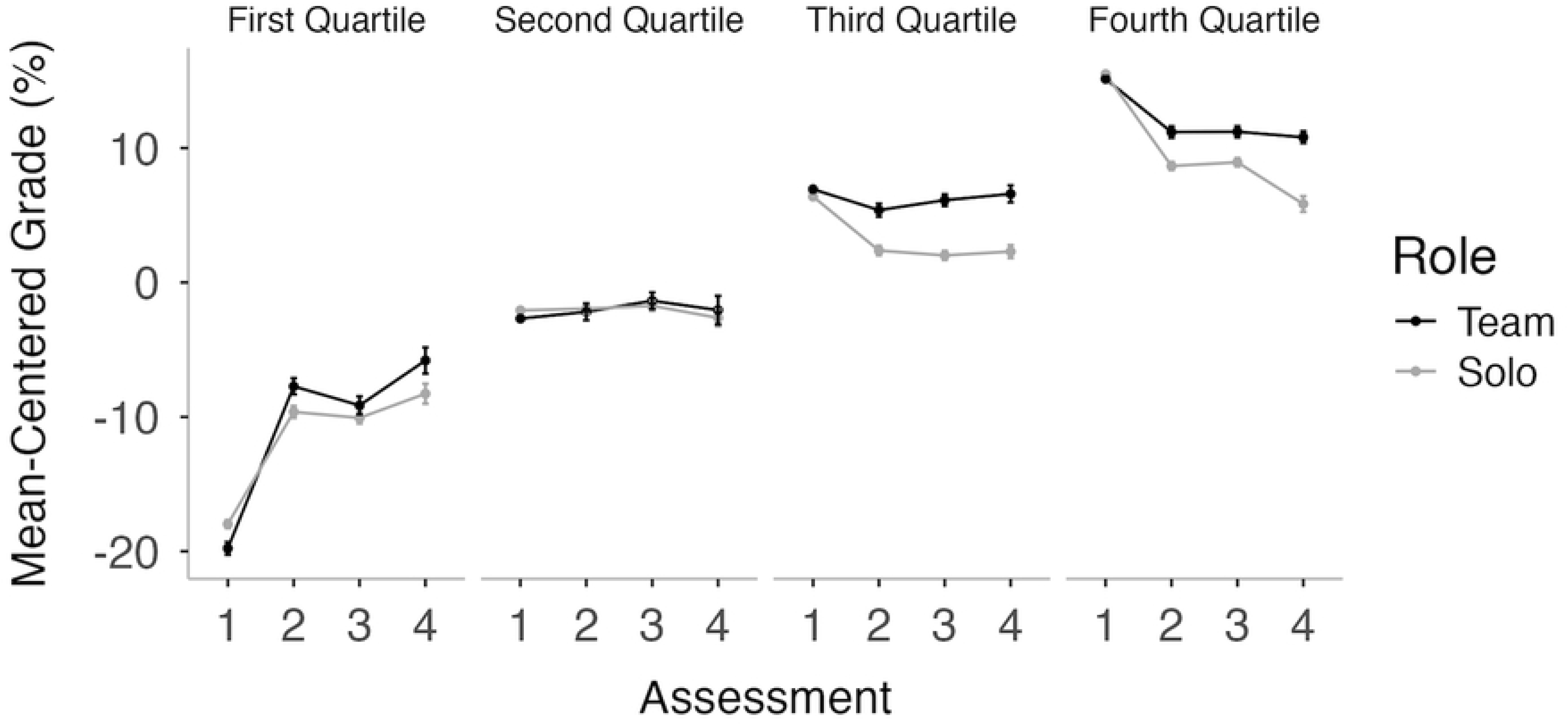
**A. Mean Scores (differences in percentage from the mean grade) in the Total Change Approach, across all assessments, averaged over all courses, separately for team and solo students.** **B Total Change Approach** Examines these differences between team and solo student outcomes as a function of how well students performed on the first assessment.

**Table S2 Total Change Approach.** Shows the mean percentage change scores for each course in the data set. The change scores were analyzed with an ANOVA model that included student role and course as fixed factors.

**Figure S5.**
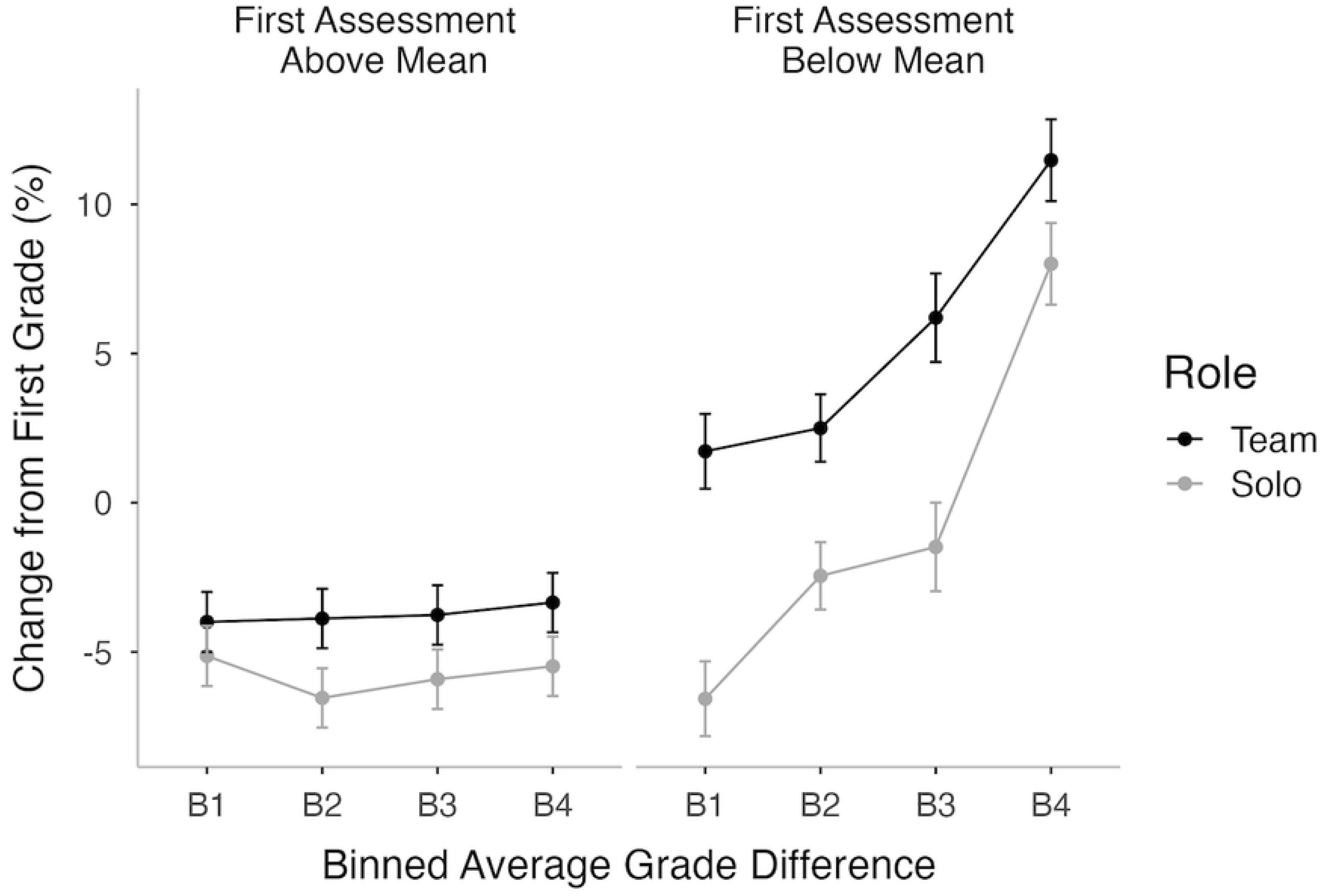
shows the average change scores following the first assessment, as a function of team heterogeneity (indexed by the standard deviation of first assessment scores in each team), separately for students scoring above versus below the mean on the first assessment, and separately for participating (team) vs non-participating (solo) students.

